# Potential mechanisms linking SIRT activity and hypoxic 2-hydroxyglutarate generation: no role for direct enzyme (de)acetylation

**DOI:** 10.1101/141416

**Authors:** Sergiy M. Nadtochiy, Yves T. Wang, Jimmy Zhang, Keith Nehrke, Xenia Schafer, Kevin Welle, Sina Ghaemmaghami, Josh Munger, Paul S. Brookes

## Abstract

2-hydroxyglutarate (2-HG) is a hypoxic metabolite with potentially important epigenetic signaling roles. The mechanisms underlying 2-HG generation are poorly understood, but evidence suggests a potential regulatory role for the sirtuin family of lysine deacetylases. Thus, we hypothesized that the acetylation status of the major 2-HG-generating enzymes (isocitrate dehydrogenase (IDH), malate dehydrogenase (MDH) and lactate dehydrogenase (LDH)) may govern their 2-HG generating activity. *In-vitro* acetylation of these enzymes, with confirmation by western blotting, mass spectrometry, and reversibility by incubation with recombinant sirtuins, yielded no effect on 2-HG generating activity. In addition, while elevated 2-HG in hypoxia is associated with the activation of lysine deacetylases, we found that mice lacking mitochondrial SIRT3 exhibited hyperacetylation and elevated 2-HG. These data suggest there is no direct link between enzyme acetylation and 2-HG production. Furthermore, our observed effects of *in-vitro* acetylation on the canonical activities of IDH, MDH and LDH appeared to contrast sharply with previous findings wherein acetyl-mimetic lysine mutations resulted in inhibition of these enzymes. Overall these data suggest that a causal relationship should not be assumed, between acetylation of metabolic enzymes and their activities, canonical or otherwise.

## INTRODUCTION

2-hydroxyglutarate (2-HG) is a non-canonical metabolite with potential signaling roles that is synthesized from the Krebs cycle metabolite α-ketoglutarate (α-KG). Among the enzymes thought to convert α-KG to 2-HG are cancer-linked forms of mitochondrial NADP+-dependent isocitrate dehydrogenase (IDH2) [1], as well as malate dehydrogenase type 2 (MDH2) [2] and lactate dehydrogenase (LDH) [3,4]. The mechanisms which trigger a neomorphic 2-HG synthetic activity are not fully elucidated.

It has been reported that a specific IDH2 mutation (R_172_K) found in glioma cells facilitates 2-HG production [1], leading to the early characterization of 2-HG as an oncometabolite. In addition, hypoxia [3–5] and acidic pH [6–8] trigger 2-HG production by MDH2 and LDH. Acidic pH is thought to protonate α-KG, enhancing its access to the substrate binding pocket of these enzymes [7]. However, it is unclear whether post-translational modification of these enzymes can also regulate 2-HG production.

Lysine acetylation, regulated by the sirtuin family of NAD+-dependent protein deacylases (SIRTs) [9], is emerging as an important regulatory mechanism for numerous metabolic processes [10–13]. Previously we demonstrated that an elevation in 2-HG levels observed in cardiac ischemic preconditioning (IPC) was associated with lysine deacetylation [14,15], and we and others showed that IPC-induced deacetylation occurs on IDH2, MDH2 and LDH-A [15–17]. Intriguingly, both lysine deacetylation and the elevation in 2-HG seen in IPC were inhibited by the SIRT1 inhibitor splitomicin (Sp) [14,15]. As such, we hypothesized herein that the 2-HG generating capacity of these dehydrogenases may be regulated by lysine acetylation status.

## METHODS

### Reagents

All chemicals and reagents were obtained from Sigma (St. Louis MO) unless otherwise stated. LDH was from bovine heart (L3916, Sigma), while MDH2 (M2634, Sigma) and NADP+ dependent mitochondrial IDH2 (I2002, Sigma) were from porcine heart. HEK293 cells were from ATCC (Manassas VA).

### In-vitro protein acetylation & western blotting

Protein acetylation was performed similar to [18]. IDH2, MDH2 or LDH (2 mg/ml) were incubated overnight at 37 °C in 50 mM Tris-HCl (pH 8) plus 1.5 mM acetyl coenzyme A, followed by neutralization of pH prior to further analysis. For MDH2, acetylated protein was further subjected to deacetylation by incubation with recombinant human SIRT3 (ab125810, AbCam, Cambridge MA).

To monitor protein acetylation, naïve and acetylated proteins were resolved on 12% SDS-PAGE gels, transferred to nitrocellulose and probed with pan anti-acetyl-lysine (K-Ac) or anti-SIRT3 antibodies (Cell Signaling, Danvers MA) at 1:1000 dilution, followed by HRP-linked secondary antibody (GE Biosciences, Pittsburgh PA) at 1:2500 and enhanced chemiluminescent detection (Pierce, Rockford IL). In addition for MDH2, acetylation at the previously published K_239_ site [17,19] was determined by mass spectrometry (Thermo Scientific Q-Exactive) in the University of Rochester Proteomics and Mass Spectrometry Core Facility. In brief, proteins were reduced and alkylated (DTT, IAA), trypsinized (Pierce) desalted (C18 Tips, Pierce), dried and suspended in 0.1% TFA. Peptides were separated by C18 HPLC (Easy nLC-1000, Thermo Fisher), with a custom column and an elution gradient of 0.1% formic acid in water → 0.1% formic acid in acetonitrile over 30 min. LC effluent was diverted to a Q Exactive Plus mass spectrometer (Thermo Fisher), operated in data-dependent mode, with a full MS1 scan (400–1400 m/z, resolution 70,000 at m/z 200, AGC target of 10^6^, max injection time 50 ms), followed by 8 data-dependent MS2 scans (resolution 35,000, AGC target of 10^5^, max injection time 120 ms). Isolation width was 1.5 m/z, offset 0.3 m/z, and normalized collision energy 27. Raw data were searched using Mascot (Matrix Science) within Proteome Discoverer software (Thermo Fisher), using the SwissProt *Sus scrofa* database with <2 missed cleavages, MS1 tolerance 10 ppm, and MS2 tolerance 25 mmu. Carbamidomethyl was set as a fixed modification, with methionine oxidation, lysine acetylation and N-term acetylation as variable modifications. Percolator was used as the FDR calculator, filtering peptides with q-value > 0.01.

### Enzyme activities and LC-MS/MS metabolite assays

Activities of IDH, MDH or LDH were measured spectrophotometrically at 340 nm, representing either the consumption of NADH (for MDH2 and LDH) or the generation of NADPH (for IDH2) [6]. In addition, aliquots from spectrophotometric cuvets were withdrawn for quantitation of selected reaction substrates or products by LC-MS/MS as previously reported [6]. <u>Note</u>: The LC-MS/MS method employed was not capable of discerning L vs. D enantiomers of 2-HG. As such we assume herein that 2-HG from IDH2 is D-2-HG, while that from LDH or MDH2 is L-2-HG [1,3,4].

### Animals & primary cells

Male littermate wild type (WT) and *Sirt3*^-/-^ mice on a C57BL/6J background were used at 2 mo. of age. All mice were maintained in a pathogen-free vivarium with a 12 hr. light/dark cycle and food and water available *ad libitum*. All procedures conformed to the National Institutes of Health Guide for the Care and Use of Laboratory Animals and were approved by the AAALAC-accredited University of Rochester Committee on Animal Resources.

For cellular hypoxia experiments, HEK293 cells were placed in a hypoxic chamber (pO_2_ <0.1 %) for 20 hr. as previously described [6]. Primary adult mouse cardiomyocytes were isolated as previously described [20]. Cytosolic pH was measured by fluorescent microscopy using the pH-sensitive probe BCECF as previously described [6]. To model ischemia, cardiomyocytes were plated on laminin-coated 25 mm glass cover slips (25,000 cells per slide) in 35 mm dishes, followed by over placement of a second cover slip to exclude oxygen and limit extracellular volume. We previously reported on a similar method using cartridges within the Seahorse XF apparatus to achieve similar ends [20,21]. Cells in the center of the cover slip were imaged after 10 min.

Mitochondria were isolated from WT or *Sirt3*^-/-^ hearts as previously described [22]. The activity of mitochondrial L-2-HG dehydrogenase was measured spectrophotometrically and by LC-MS/MS as previously described [6].

### Statistics

Significance between groups was determined by ANOVA, assuming a normal distribution, followed by post-hoc Student’s t-test. Differences were considered statistically significant at p<0.05, and graphs show means ± standard errors.

## RESULTS

We previously demonstrated a role for SIRT1 in the elevation of 2-HG observed in cardiac ischemic preconditioning (IPC) [14]. In addition, 2-HG is elevated in cells exposed to hypoxia [3,4]. However, since these are distinct experimental systems (intermittent whole organ ischemia vs. prolonged cellular hypoxia), it is not clear if SIRT1 is also required for the latter phenomenon. To investigate this we measured 2-HG levels in HEK293 cells exposed to hypoxia with or without the SIRT1 inhibitor splitomicin (Sp). As expected, hypoxia markedly increased 2-HG levels (Figure 1A), and notably this effect was blunted by Sp, suggesting a similar role for SIRT1 in both hypoxic situations.

**Figure 1.**
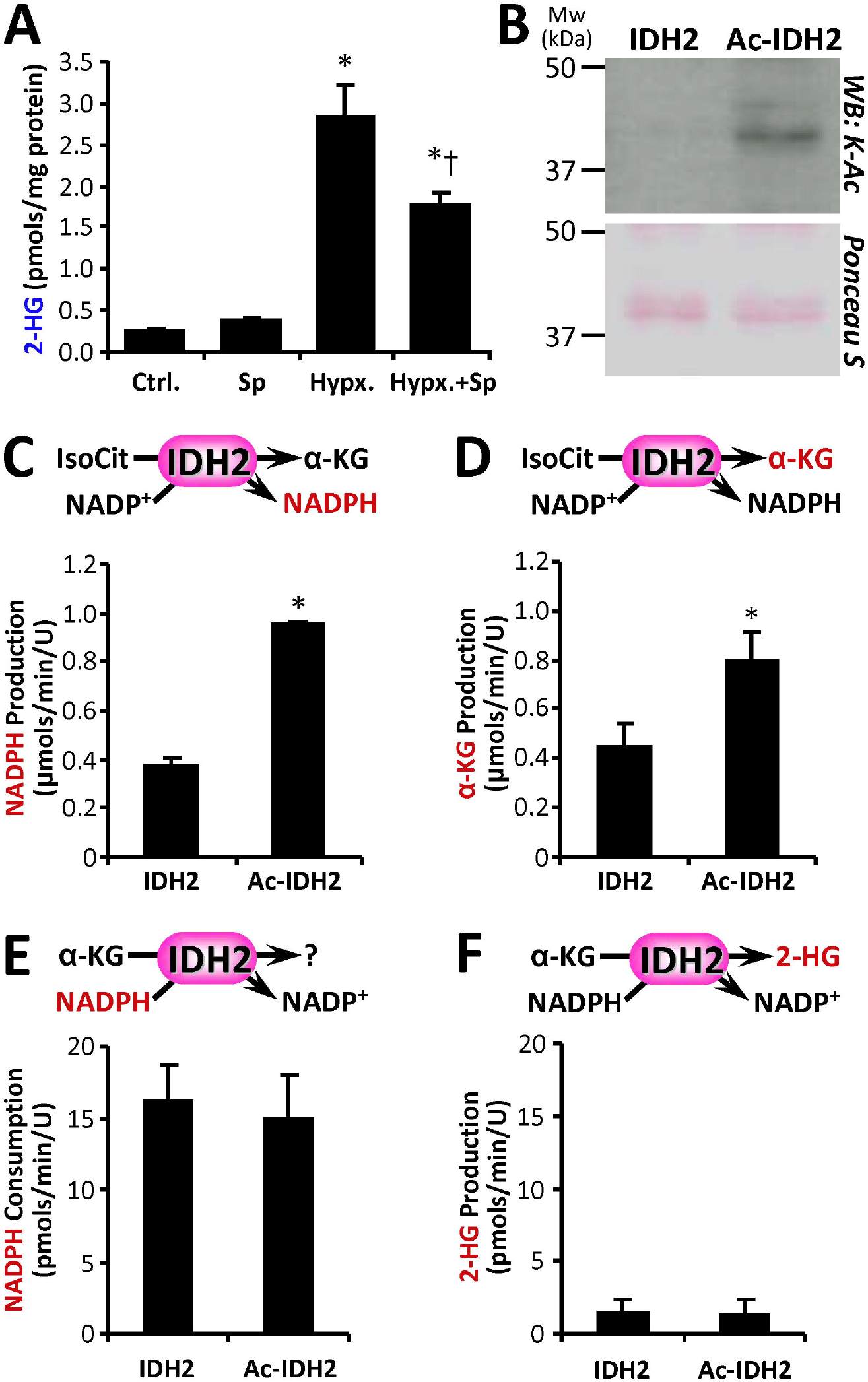
Cell Hypoxia, IDH2 Acetylation & Activities. **(A)**: HEK293 cells were exposed to hypoxia with or without the SIRT1 inhibitor splitomicin (Sp), followed by measurement of 2-HG levels. Data are means ± SEM, n=5. *p<0.05 vs. Ctrl. †p<0.05 vs. Hypoxia alone. **(B)**: Purified IDH2 was acetylated *in-vitro* with acetyl-CoA (see methods), followed by western blot probe with anti acetyl-lysine (K-Ac) antibody. Ponceau S stained membrane below indicates protein loading. Images representative of at least 4 independent experiments. **(C/D)**: Canonical activity of naïve and acetylated IDH2 with isocitrate and NADP+ as substrates was measured spectrophotometrically as NADPH production (C) or by LC-MS/MS as α-KG production (D). **(E/F)**: Non-canonical reductive activity of IDH2 with α-KG and NADPH as substrates was measured spectrophotometrically as NADPH consumption (E) or by LC-MS/MS as 2-HG production (F). All enzyme rate data are means ± SEM, n=4. *p<0.05 between naïve and Ac-IDH2. Reactions monitored are shown above each graph, with the metabolite measured shown in red font.

IDH2, MDH2 and LDH are enzymes capable of generating 2-HG [1,3,4,6,7] and are also thought to be targets for SIRT mediated deacetylation [15–17,19,23,24]. Thus, we investigated whether the acetylation status of these enzymes affected their 2-HG generating capacity. Purified NADP^+^-dependent mitochondrial IDH2 was acetylated *in-vitro* (Figure 1B), and its canonical reaction (i.e., NADP+ + isocitrate → α-KG + NADPH) was monitored both spectrophotometrically and by LC-MS/MS [6]. Acetylated IDH2 (Ac-IDH2) exhibited significantly higher activity than the naïve enzyme, measured either as NADPH or α-KG generation (Figures 1C and 1D respectively). This result contrasts with a previous study [25], wherein an acetyl-mimetic IDH2 mutation (K_413_Q) demonstrated lower activity.

In terms of 2-HG generation, Figures 1E and 1F show that both naïve and Ac-IDH2 could use α-KG + NADPH as substrates, but neither generated substantial corresponding amounts of 2-HG. Instead, IDH2 is known to operate in reverse, using α-KG in a reductive carboxylation reaction to produce isocitrate [6,26]. Together these data suggest that acetylation boosted the canonical activity of IDH2 (i.e., isocitrate → α-KG), with no effect on its non-canonical activities (2-HG generation or reductive carboxylation). The fact that acetylation *did* impact IDH2 canonical activity suggests the lack of an effect on non-canonical activities (negative result) was not simply due to confounding experimental issues.

The mitochondrial enzyme MDH2 can also produce 2-HG [3,4,6,7], and notably lysine 239 of MDH2 is differentially acetylated in mice lacking the mitochondrial deacetylase SIRT3 [17]. Furthermore, an acetyl-mimetic K_239_Q mutant protein exhibits lower MDH activity [17], suggesting acetylation at this site is inhibitory. We acetylated purified MDH2 *in-vitro* [18] and verified this by western blot. As shown in Figure 2A this acetylation event was also reversed by incubation with human recombinant SIRT3 + NAD+. In addition, mass spectrometric analysis of naïve vs. acetylated MDH2 (Ac-MDH2) (Figure 2B–D) revealed a difference of 42 mass units in the y^1+^ ion for the peptide containing K_239_ (IQEAGTEVVK underlined in panel 2C), confirming acetylation at this site.

**Figure 2.**
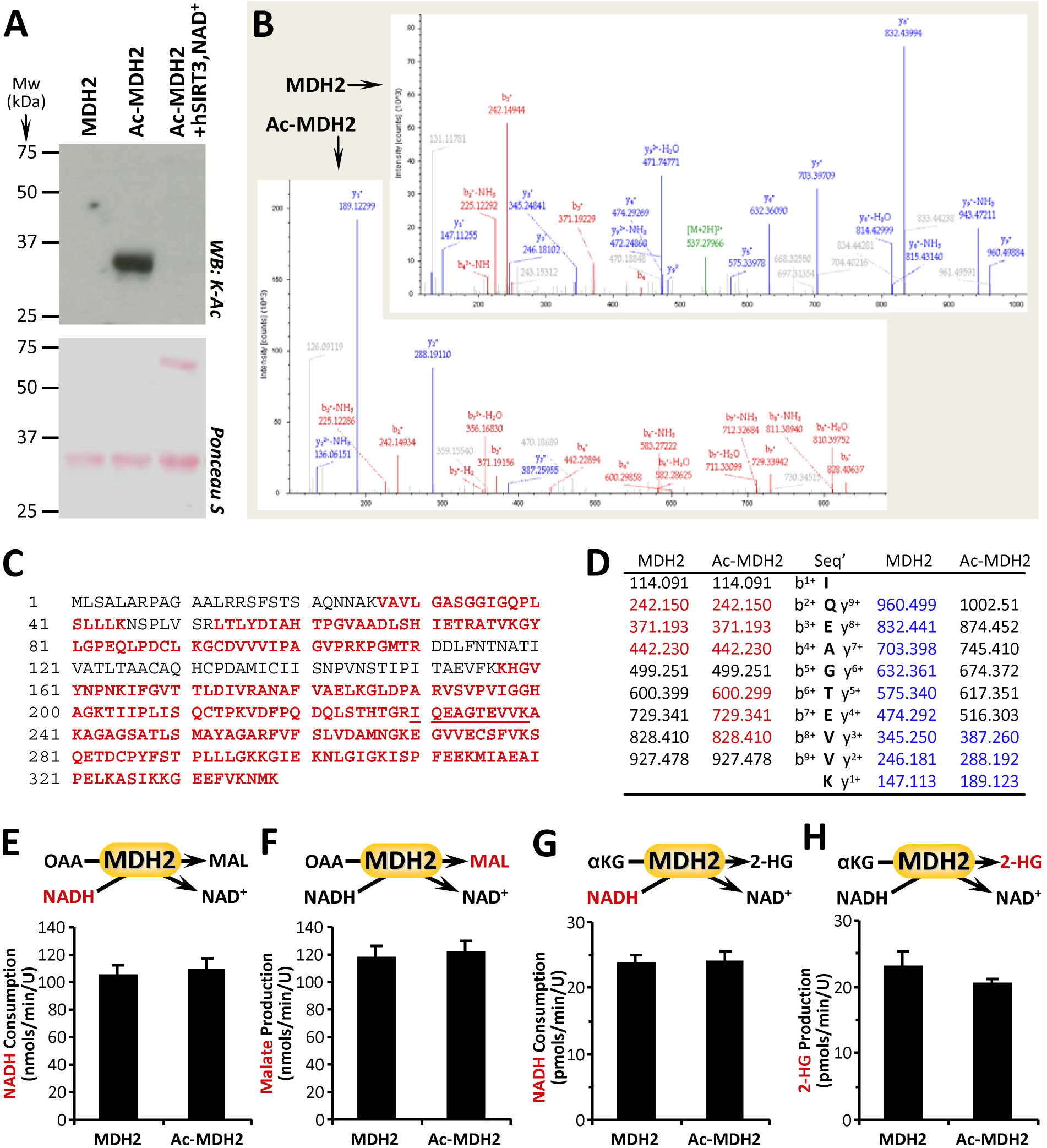
MDH2 Acetylation & Activities. **(A)**: Purified MDH2 was acetylated *in-vitro* with acetyl-CoA (see methods), followed by western blot probe with anti acetyl-lysine (K-Ac) antibody. Ponceau S stained membrane below indicates protein loading. Images representative of at least 4 independent experiments. **(B/C/D)**: Naïve and acetylated MDH2 (Ac-MDH2) were digested with trypsin and resultant peptides analyzed by mass spectrometry to identify acetylation sites. 76% sequence coverage was obtained (panel C, red font) including the peptide containing the proposed K_239_ acetylation site (underlined). Sample spectra are shown in B, with panel D listing identified peptides corresponding to b^n+^ ions (red) and y^n+^ ions (blue). A difference of 42 mass units between y^+1^ ion in MDH2 vs. Ac-MDH2 (189.123 – 147.113) indicates acetylation. **(E/F)**: Canonical activity of naïve and acetylated MDH2 with OAA and NADH as substrates was measured spectrophotometrically as NADH consumption (E) or by LC-MS/MS as malate production (F). **(G/H)**: Non-canonical activity of MDH2 with α-KG and NADH as substrates was measured spectrophotometrically as NADH consumption (G) or by LC-MS/MS as 2-HG production (H). All enzyme rate data are means ± SEM, n=4–6. Reactions monitored are shown above each graph, with the metabolite measured shown in red font.

Surprisingly, as shown in Figures 2E and 2F, the canonical MDH enzymatic reaction (i.e., OAA + NADH → malate + NAD+) was no different between naïve vs. Ac-MDH2 (Note: the normal forward reaction, malate → OAA, is energetically unfavorable *in-vitro* so the enzyme is routinely assayed in the reverse direction). Furthermore, with α-KG as a substrate the enzymatic reaction to generate 2-HG was also unaffected by acetylation status (Figures 2G and 2H). Overall, these data suggest that neither canonical nor alternative activities of MDH2 were impacted by acetylation.

Another enzyme implicated in 2-HG generation is LDH, and notably lysine 5 within LDH-A was identified as an acetylation site [27], with acetyl-mimetic mutation of this residue (K5Q) resulting in enzyme inhibition. In contrast to these mutant enzyme findings, when we acetylated purified LDH *in-vitro* (Figure 3A), we observed no effect on canonical activity (pyruvate + NADH → lactate + NAD+) (Figures 3B and 3C). Furthermore, 2-HG generation from α-KG by LDH was not affected by acetylation (Figures 3D and 3E). Together these data suggest that acetylation does not impact the enzymatic properties of LDH.

**Figure 3.**
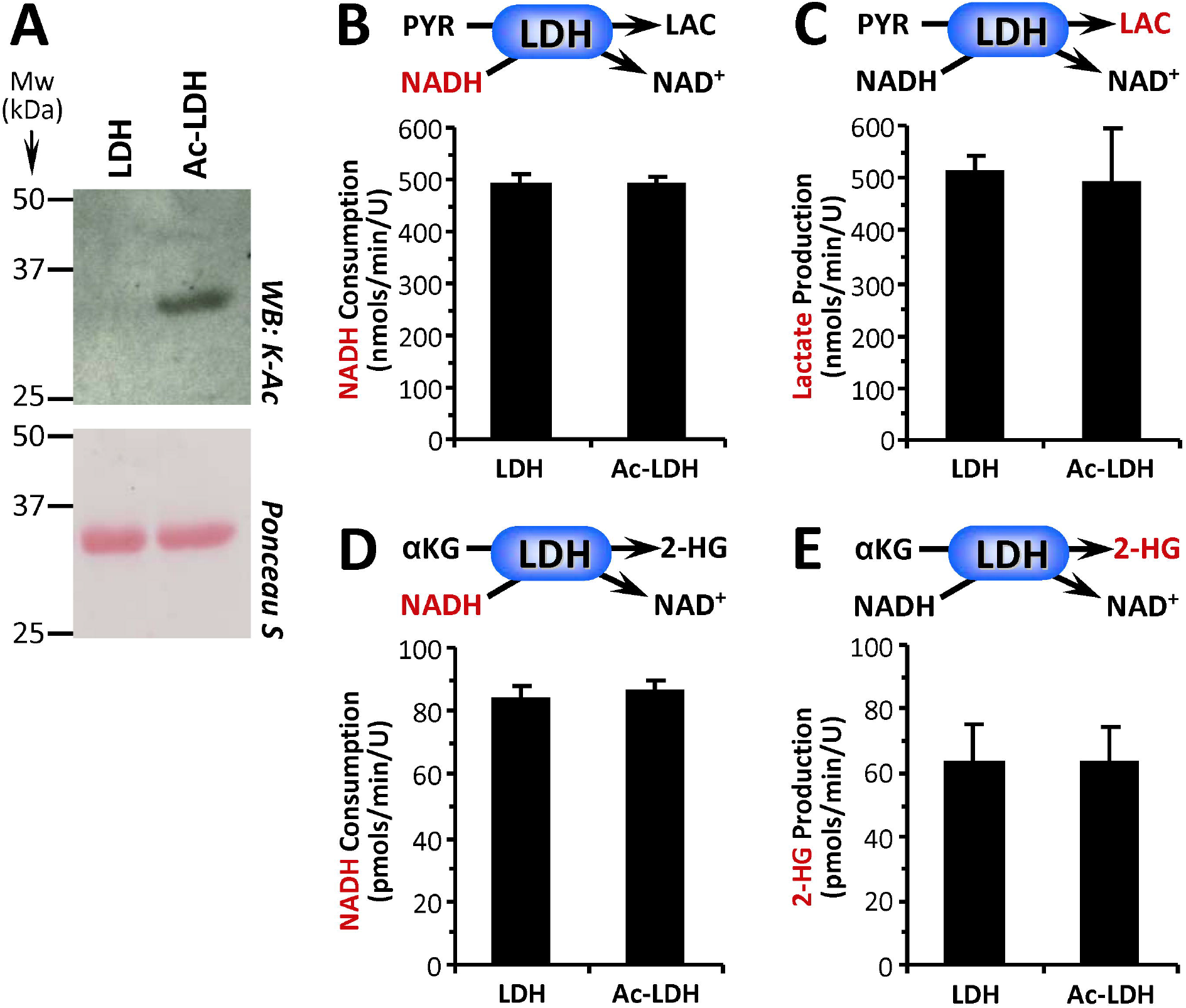
LDH Acetylation & Activities. **(A)**: Purified LDH was acetylated *in-vitro* with acetyl-CoA (see methods), followed by western blot probe with anti acetyl-lysine (K-Ac) antibody. Ponceau S stained membrane below indicates protein loading. Images representative of at least 4 independent experiments. **(B/C)**: Canonical activity of naïve and acetylated LDH with pyruvate and NADH as substrates was measured spectrophotometrically as NADH consumption (B) or by LC-MS/MS as lactate production (C). **(D/E)**: Non-canonical activity of LDH with-KG and NADH as substrates was measured spectrophotometrically as NADH consumption (D) or by LC-MS/MS as 2-HG production (E). All enzyme rate data are means ± SEM, n=4–7. Reactions monitored are shown above each graph, with the metabolite measured shown in red font.

To further investigate the possible role of lysine acetylation as a regulator of 2-HG generation, we examined mice with deletion of *Sirt3* (Figure 4A), a mitochondrial SIRT known to deacetylate IDH2, MDH2 and LDH [15–17,19,23,24]. As shown in Figures 4B and 4C, *Sirt3*^-/-^ hearts exhibited the expected hyper-acetylation of cardiac mitochondrial proteins (vs. wild-type, WT), but surprisingly this was accompanied by a 3.5-fold elevation in 2-HG. Such a result contrasts with the situation in both IPC [14] and cellular anoxia (Figure 1A), wherein deacetylation is associated with elevated 2-HG. Together these data support a disconnect between acetylation and 2-HG generation.

**Figure 4.**
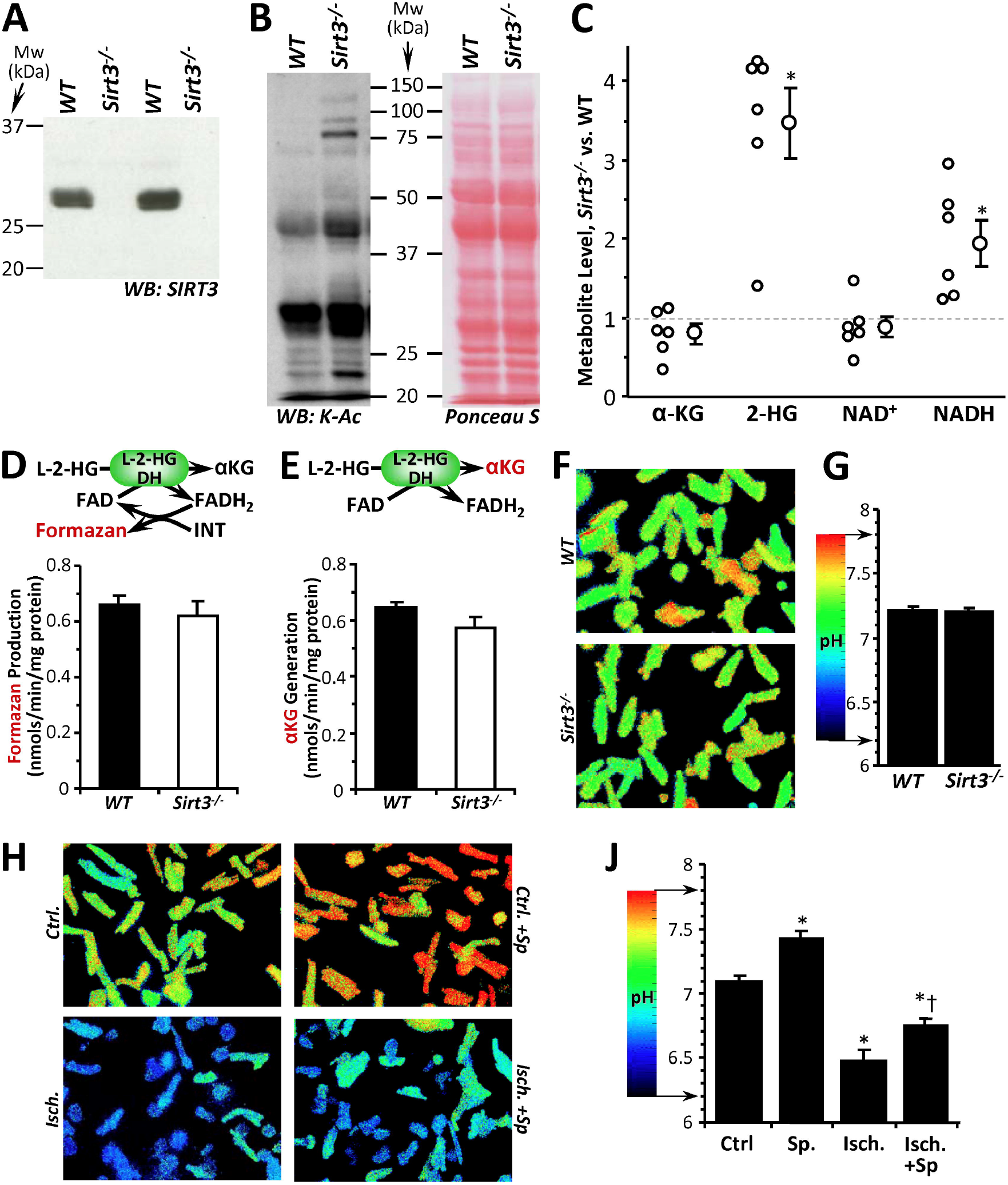
Acetylation & 2-HG in *Sirt3*^-/-^ mice, and cytosolic pH. **(A)**: Western blot shows absence of SIRT3 protein in cardiac tissue from *Sirt3*^-/-^ mice (vs. WT). **(B)**: Western blot shows protein acetylation (pan acetyl-lysine antibody) in cardiac mitochondria from WT and *Sirt3*^-/-^ mice. Ponceau S stained membrane indicates protein loading. Blots are representative of at least 4 independent experiments. **(C)**: LC-MS/MS based metabolite profiling of WT vs. *Sirt3*^-/-^ cardiac tissue. Data are expressed as the metabolite level in *Sirt3*^-/-^ relative to paired WT samples, with individual data points (n=6 pairs) alongside means ± SEM. *p<0.05 between WT and *Sirt3*^-/-^. **(D/E)**: L-2-HGDH activity in cardiac mitochondria from WT and *Sirt3*^-/-^ mice was measured spectrophotometrically via formazan-linked assay (D) or by LC-MS/MS as α-KG formation (E). Enzyme rate data are means ± SEM, n=5. Reactions monitored are shown above each graph, with the metabolite measured shown in red font. **(F/G)**: Cytosolic pH measurements in primary cultured adult cardiomyocytes from WT and *Sirt3*^-/-^ mice. Alongside representative images (F), graph (G) shows quantitation of pH (means ± SEM) from 4 independent cell preparations. **(H/J)**: Cytosolic pH measurements in primary cultured adult cardiomyocytes from WT mice subjected to simulated ischemia (see methods), with or without the SIRT1 inhibitor splitomicin (Sp). Alongside representative images (H), graph (J) shows quantitation of pH (means ± SEM) from 4–6 independent cell preparations. *p<0.05 vs. Ctrl. †p<0.05 vs. Ischemia alone.

Nevertheless, these data raise an intriguing question as to the mechanism of 2-HG elevation in *Sirt3*^-/-^ mice. One possibility may be inhibition of the mitochondrial 2-HG catabolic enzyme L-2-HG dehydrogenase (L-2-HGDH) [28]. However, as shown in Figures 4D and 4E, we observed no difference in L-2-HGDH activity between *Sirt3*^-/-^ and WT cardiac mitochondria.

Another potentially important regulator of 2-HG generation is acidic pH [6,7]. However, we did not observe any significant difference in pH between cardiomyocytes isolated from WT or *Sirt3*^-/-^ hearts (Figures 4F and 4G). Notably (and in agreement with previous studies), *Sirt3*^-/-^ hearts exhibited elevated NADH levels (Figure 4C), a phenomenon that has been attributed to inhibition of mitochondrial respiratory complex I (NADH ubiquinone oxidoreductase) [22,29]. It is thought that excessive NADH accumulation (a.k.a. *reductive stress)*, is a key driver of the α-KG → 2-HG reaction [30], and together these findings suggest that a likely cause of elevated 2-HG in *Sirt3*^-/-^ hearts is increased drive from NADH.

Finally, to uncover the basis for Sp sensitivity in the hypoxic increase in 2-HG ([14] and Figure 1A), we measured the effects of Sp on ischemia-induced acidification of primary mouse cardiomyocytes. As shown in Figures 4H and 4J, Sp elicited a slight alkalinization alone and blunted the acidification caused by ischemia. These data suggest that the role of SIRT1 in the elevation of 2-HG in hypoxia/ischemia, is likely at a position upstream of acidification.

## DISCUSSION & CONCLUSIONS

The conclusions of this study are as follows: (i) IDH2 (either naïve or acetylated) did not produce 2-HG; although Ac-IDH2 showed higher canonical activity (isocitrate → α-KG); (ii) both canonical enzyme activities and 2-HG generation by Ac-MDH2 and Ac-LDH were similar to the corresponding naïve enzymes; (iii) 2-HG was elevated in *Sirt3*^-/-^ hearts, concurrent with higher NADH levels; and (iv) the ability of the SIRT1 inhibitor splitomicin to blunt 2-HG elevation in hypoxia/ischemia may be due to an upstream role for SIRT1 in regulating acidosis. Overall, these data suggest that direct acetylation of the major identified 2-HG generating enzymes is not a mechanism that regulates 2-HG.

Our observation that IDH2 acetylation by a physiologically relevant mechanism (acetyl-CoA) increases its canonical activity (Figure 1C and 1D) is an interesting result. Since acetyl-CoA is the upstream substrate for the Krebs cycle, it is possible that acetyl-CoA mediated IDH2 acetylation represents a novel feed-forward activation mechanism for the cycle. The importance of this relative to the already well-characterized regulation of this cycle by numerous allosteric mechanisms (e.g., Ca^2^+, NADH, phosphorylation) [31–33] remains to be seen.

A consistent finding in the current study was a sharp contrast with previous observations using acetyl-mimetic mutations in dehydrogenase enzymes [17,25,27]. Specifically, acetyl-mimetic K_413_Q of IDH2, K_239_Q of MDH2 and K_5_Q of LDH-A were all shown to result in lower activities for these enzymes. In contrast we found that acetylation stimulated IDH2 activity, while having no effect on MDH2 or LDH. The acetylation methods used herein were physiologically relevant [18], and acetylation was verified by western blot, by mass spectrometry and by reversibility with a recombinant SIRT. Interpreting mutational studies can be problematic when mutations result in loss of function, and in the lysine acetylation field there are vanishingly few examples of acetyl-mimetic mutation resulting in stimulation of activity. As such, we speculate that the results of some mutational studies reporting on enzyme inhibition may be artifactual, due to unexpected effects of mutations on protein folding or other enzyme properties. Indeed, it was shown that the acetyl-mimetic mutants K_3_Q and K_118_Q of MDH isoform 1 had no effect on activity, while deacetylation mimetics K_3_R and K_118_R were inhibitory [34]. Thus, in particular for MDH, the relationship between acetylation and enzyme activity is not clear.

An important caveat in interpreting our findings, is subcellular compartmentation. In cardiomyocytes SIRT1 is cytosolic [15,35], and as shown in Figure 4 the SIRT1 inhibitor Sp results in alkalinization of the cardiomyocyte cytosol. Sp also blunts the hypoxic increase in 2-HG (Figure 1A). However, acetylation of cytosolic LDH does not impact 2-HG generation. Thus we conclude that SIRT1 may play a role in regulating 2-HG at a point upstream of acidification. In contrast, SIRT3 is mitochondrial and its genetic ablation has no effect on cytosolic pH, and no effect on L-2HGDH activity. Furthermore, no effect of acetylation on 2-HG generation was seen for the mitochondrial enzymes MDH2 or IDH2. Thus, in the case of *Sirt3*^-/-^, we speculate that elevated 2-HG originates at the level of reductive stress (excess NADH), possibly due to inhibition of mitochondrial complex I [22,29]. Interestingly, the *Sirt3*^-/-^ mouse exhibits acetylation of cytosolic LDH [17], and we showed that SIRT1 inhibition resulted in acetylation of numerous mitochondrial enzymes [15]. Thus, there appear to be signaling mechanisms by which inactivation (genetic or pharmacologic) of a SIRT in one cell compartment can impact acetylation in another compartment. Overall, regardless the compartment, it is clear from our results that direct regulation of acetylation of 2-HG generating enzymes by SIRT1 or SIRT3 does not impact 2-HG levels.

It was recently proposed that conversion of α-KG to 2-HG is a quantitatively significant mechanism by which “cells handle a mounting pool of reducing equivalents” such as excess NADH [30]. 2-HG was also proposed to represent a “redox reservoir”, to enable cells to handle reductive stress [30]. However, the canonical activities of LDH and MDH2 are at least 3-orders of magnitude greater than the rates at which these enzymes reduce α-KG to 2-HG [6,2]. In conditions of hypoxia and reductive stress, the 2-HG reaction is quantitatively insignificant as a consumer of NADH, relative to the role of LDH (pyruvate + NADH → lactate + NAD+). In hypoxia, 2-HG generation is also several orders of magnitude smaller than another important redox sink, namely the generation of succinate from fumarate by reversal of respiratory complex II [36–38]. Furthermore, 2-HG is not a true redox reservoir because the reverse reaction to regenerate α-KG from 2-HG (i.e., the reaction catalyzed by L-2-HGDH) uses FAD as an electron acceptor and is not known to regenerate NADH.

These quantitative limitations on the role of 2-HG as a redox sink do not diminish the potential importance of 2-HG as a hypoxic signaling molecule [6,39,40]. In addition, the role of other protein post-translational modifications that may regulate 2-HG under conditions such as hypoxia and/or differential acetylation is worthy of further investigation.

## ACKNOWLEDGEMENTS

This work was funded by a grant from the US National Institutes of Health (RO1 HL-071158) to PSB. We thank Dr. George Porter Jr. for *Sirt3*^-/-^ mice.

## REFERENCES

1 Gross, S., Cairns, R. A., Minden, M. D., Driggers, E. M., Bittinger, M. A., Jang, H. G., Sasaki, M., Jin, S., Schenkein, D. P., Su, S. M., et al. (2010) Cancer-associated metabolite 2-hydroxyglutarate accumulates in acute myelogenous leukemia with isocitrate dehydrogenase 1 and 2 mutations. J. Exp. Med. 207, 339–344.

2 Rzem, R., Vincent, M.-F., Van Schaftingen, E. and Veiga-da-Cunha, M. (2007) l-2-Hydroxyglutaric aciduria, a defect of metabolite repair. J. Inherit. Metab. Dis. 30, 681–689.

3 Intlekofer, A. M., Dematteo, R. G., Venneti, S., Finley, L. W. S., Lu, C., Judkins, A. R., Rustenburg, A. S., Grinaway, P. B., Chodera, J. D., Cross, J. R., et al. (2015) Hypoxia Induces Production of L-2-Hydroxyglutarate. Cell Metab. 22, 304–311.

4 Oldham, W. M., Clish, C. B., Yang, Y. and Loscalzo, J. (2015) Hypoxia-Mediated Increases in l-2-hydroxyglutarate Coordinate the Metabolic Response to Reductive Stress. Cell Metab. 22, 291–303.

5 Armitage, E. G., Kotze, H. L., Allwood, J. W., Dunn, W. B., Goodacre, R. and Williams, K. J. (2015) Metabolic profiling reveals potential metabolic markers associated with Hypoxia Inducible Factor-mediated signalling in hypoxic cancer cells. Sci. Rep. 5, 15649.

6 Nadtochiy, S. M., Schafer, X., Fu, D., Nehrke, K., Munger, J. and Brookes, P. S. (2016) Acidic pH Is a Metabolic Switch for 2-Hydroxyglutarate Generation and Signaling. J. Biol. Chem. 291, 20188–20197.

7 Intlekofer, A. M., Wang, B., Liu, H., Shah, H., Carmona-Fontaine, C., Rustenburg, A. S., Salah, S., Gunner, M. R., Chodera, J. D., Cross, J. R., et al. (2017) L-2-Hydroxyglutarate production arises from noncanonical enzyme function at acidic pH. Nat. Chem. Biol. 13, 494–500.

8 Teng, X., Emmett, M. J., Lazar, M. A., Goldberg, E. and Rabinowitz, J. D. (2016) Lactate Dehydrogenase C Produces S-2-Hydroxyglutarate in Mouse Testis. ACS Chem. Biol. 11, 2420–7.

9 Anderson, K. A., Green, M. F., Huynh, F. K., Wagner, G. R. and Hirschey, M. D. (2014) SnapShot: Mammalian Sirtuins. Cell 159, 956–956.e1.

10 Huang, J.-Y., Hirschey, M. D., Shimazu, T., Ho, L. and Verdin, E. (2010) Mitochondrial sirtuins. Biochim. Biophys. Acta - Proteins Proteomics 1804, 1645–1651.

11 Correia, M., Perestrelo, T., Rodrigues, A. S., Ribeiro, M. F., Pereira, S. L., Sousa, M. I. and Ramalho-Santos, J. (2017) Sirtuins in metabolism, stemness and differentiation. Biochim. Biophys. Acta - Gen. Subj. 1861, 3444–3455.

12 van de Ven, R. A. H., Santos, D. and Haigis, M. C. (2017) Mitochondrial Sirtuins and Molecular Mechanisms of Aging. Trends Mol. Med. 23, 320–331.

13 Ye, X., Li, M., Hou, T., Gao, T., Zhu, W. and Yang, Y. (2015) Sirtuins in glucose and lipid metabolism. Oncotarget 8, 1845–1859.

14 Nadtochiy, S. M., Urciuoli, W., Zhang, J., Schafer, X., Munger, J. and Brookes, P. S. (2015) Metabolomic profiling of the heart during acute ischemic preconditioning reveals a role for SIRT1 in rapid cardioprotective metabolic adaptation. J. Mol. Cell. Cardiol. 88, 64–72.

15 Nadtochiy, S. M., Redman, E., Rahman, I. and Brookes, P. S. (2011) Lysine deacetylation in ischaemic preconditioning: the role of SIRT1. Cardiovasc. Res. 89, 643–649.

16 Parker, B. L., Shepherd, N. E., Trefely, S., Hoffman, N. J., White, M. Y., Engholm-Keller, K., Hambly, B. D., Larsen, M. R., James, D. E. and Cordwell, S. J. (2014) Structural basis for phosphorylation and lysine acetylation cross-talk in a kinase motif associated with myocardial ischemia and cardioprotection. J. Biol. Chem. 289, 25890–906.

17 Hebert, A. S., Dittenhafer-Reed, K. E., Yu, W., Bailey, D. J., Selen, E. S., Boersma, M. D., Carson, J. J., Tonelli, M., Balloon, A. J., Higbee, A. J., et al. (2013) Calorie restriction and SIRT3 trigger global reprogramming of the mitochondrial protein acetylome. Mol. Cell 49, 186–99.

18 Wagner, G. R. and Payne, R. M. (2013) Widespread and enzyme-independent Nε-acetylation and Nε-succinylation of proteins in the chemical conditions of the mitochondrial matrix. J. Biol. Chem. 288, 29036–45.

19 Still, A. J., Floyd, B. J., Hebert, A. S., Bingman, C. A., Carson, J. J., Gunderson, D. R., Dolan, B. K., Grimsrud, P. A., Dittenhafer-Reed, K. E., Stapleton, D. S., et al. (2013) Quantification of mitochondrial acetylation dynamics highlights prominent sites of metabolic regulation. J. Biol. Chem. 288, 26209–19.

20 Nadtochiy, S. M., Madukwe, J., Hagen, F. and Brookes, P. S. (2014) Mitochondrially targeted nitro-linoleate: a new tool for the study of cardioprotection. Br. J. Pharmacol. 171, 2091–2098.

21 Guo, S., Olm-Shipman, A., Walters, A., Urciuoli, W. R., Devito, S., Nadtochiy, S. M., Wojtovich, A. P. and Brookes, P. S. (2012) A cell-based phenotypic assay to identify cardioprotective agents. Circ. Res. 110, 948–57.

22 Porter, G. A., Urciuoli, W. R., Brookes, P. S. and Nadtochiy, S. M. (2014) SIRT3 deficiency exacerbates ischemia-reperfusion injury: implication for aged hearts. Am. J. Physiol. Heart 306, 1602–9.

23 Sol, E. M., Wagner, S. A., Weinert, B. T., Kumar, A., Kim, H.-S., Deng, C.-X. and Choudhary, C. (2012) Proteomic investigations of lysine acetylation identify diverse substrates of mitochondrial deacetylase sirt3. PLoS One 7, e50545.

24 Rardin, M. J., Newman, J. C., Held, J. M., Cusack, M. P., Sorensen, D. J., Li, B., Schilling, B., Mooney, S. D., Kahn, C. R., Verdin, E., et al. (2013) Label-free quantitative proteomics of the lysine acetylome in mitochondria identifies substrates of SIRT3 in metabolic pathways. Proc. Natl. Acad. Sci. U. S. A. 110, 6601–6.

25 Yu, W., Dittenhafer-Reed, K. E. and Denu, J. M. (2012) SIRT3 Protein Deacetylates Isocitrate Dehydrogenase 2 (IDH2) and Regulates Mitochondrial Redox Status. J. Biol. Chem. 287, 14078–14086.

26 Wise, D. R., Ward, P. S., Shay, J. E. S., Cross, J. R., Gruber, J. J., Sachdeva, U. M., Platt, J. M., DeMatteo, R. G., Simon, M. C. and Thompson, C. B. (2011) Hypoxia promotes isocitrate dehydrogenase-dependent carboxylation of α-ketoglutarate to citrate to support cell growth and viability. Proc. Natl. Acad. Sci. U. S. A. 108, 19611–6.

27 Zhao, D., Zou, S.-W., Liu, Y., Zhou, X., Mo, Y., Wang, P., Xu, Y.-H., Dong, B., Xiong, Y., Lei, Q.-Y., et al. (2013) Lysine-5 Acetylation Negatively Regulates Lactate Dehydrogenase A and Is Decreased in Pancreatic Cancer. Cancer Cell 23, 464–476.

28 Rzem, R., Achouri, Y., Marbaix, E., Schakman, O., Wiame, E., Marie, S., Gailly, P., Vincent, M.-F., Veiga-da-Cunha, M. and Van Schaftingen, E. (2015) A Mouse Model of L-2-Hydroxyglutaric Aciduria, a Disorder of Metabolite Repair. PLoS One 10, e0119540.

29 Ahn, B.-H., Kim, H.-S., Song, S., Lee, I. H., Liu, J., Vassilopoulos, A., Deng, C.-X. and Finkel, T. (2008) A role for the mitochondrial deacetylase Sirt3 in regulating energy homeostasis. Proc. Natl. Acad. Sci. U. S. A. 105, 14447–52.

30 Loscalzo, J. (2016) Adaptions to Hypoxia and Redox Stress: Essential Concepts Confounded by Misleading Terminology. Circ. Res. 119, 511–3.

31 Hofmeyr, J. H. S., Rohwer, J. M. and Snoep, J. L. (2006) Conditions for effective allosteric feedforward and feedback in metabolic pathways. Syst. Biol. 153, 327–31.

32 Bouskila, M., Hunter, R. W., Ibrahim, A. F. M., Delattre, L., Peggie, M., van Diepen, J. A., Voshol, P. J., Jensen, J. and Sakamoto, K. (2010) Allosteric Regulation of Glycogen Synthase Controls Glycogen Synthesis in Muscle. Cell Metab. 12, 456–466.

33 Tanaka, T., Sue, F. and Morimura, H. (1967) Feed-forward activation and feed-back inhibition of pyruvate kinase type L of rat liver. Biochem. Biophys. Res. Commun. 29, 444–449.

34 Kim, E. Y., Kim, W. K., Kang, H. J., Kim, J.-H., Chung, S. J., Seo, Y. S., Park, S. G., Lee, S. C. and Bae, K.-H. (2012) Acetylation of malate dehydrogenase 1 promotes adipogenic differentiation via activating its enzymatic activity. J. Lipid Res. 53, 1864–1876.

35 Tanno, M., Sakamoto, J., Miura, T., Shimamoto, K. and Horio, Y. (2007) Nucleocytoplasmic shuttling of the NAD+-dependent histone deacetylase SIRT1. J. Biol. Chem. 282, 6823–32.

36 Chouchani, E. T., Pell, V. R., Gaude, E., Aksentijević, D., Sundier, S. Y., Robb, E. L., Logan, A., Nadtochiy, S. M., Ord, E. N. J., Smith, A. C., et al. (2014) Ischaemic accumulation of succinate controls reperfusion injury through mitochondrial ROS. Nature 515, 431–435.

37 Sanadi, D. R. and Fluharty, A. L. On The Mechanism Of Oxidative Phosphorylation. Vii. The Energy-Requiring Reduction Of Pyridine Nucleotide By Succinate And The Energy-Yielding Oxidation Of Reduced Pyridine Nucleotide By Fumarate. Biochemistry 2, 523–8.

38 Hochachka, P. W., Owen, T. G., Allen, J. F. and Whittow, G. C. (1975) Multiple end products of anaerobiosis in diving vertebrates. Comp. Biochem. Physiol. B. 50, 17–22.

39 Su, X., Wellen, K. E. and Rabinowitz, J. D. (2016) Metabolic control of methylation and acetylation. Curr. Opin. Chem. Biol. 30, 52–60.

40 Xu, W., Yang, H., Liu, Y., Yang, Y., Wang, P., Kim, S.-H., Ito, S., Yang, C., Wang, P., Xiao, M.-T., et al. (2011) Oncometabolite 2-Hydroxyglutarate Is a Competitive Inhibitor of α-Ketoglutarate-Dependent Dioxygenases. Cancer Cell 19, 17–30.

